# Improved temporal resolution for mapping brain metabolism using functional PET and anatomical MRI knowledge

**DOI:** 10.1101/2020.07.08.192872

**Authors:** Viswanath P. Sudarshan, Shenpeng Li, Sharna D. Jamadar, Gary F. Egan, Suyash P. Awate, Zhaolin Chen

## Abstract

Functional positron emission tomography (fPET) imaging using continuous infusion of [18F]-fluorodeoxyglucose (FDG) is a novel neuroimaging technique to track dynamic glucose utilization in the brain. In comparison to conventional static PET, fPET maintains a sustained supply of glucose in the blood plasma which improves sensitivity to measure dynamic glucose changes in the brain, and enables mapping of dynamic brain activity in task-based and resting-state fPET studies. However, there is a trade-off between temporal resolution and spatial noise due to the low concentration of FDG and the limited sensitivity of multi-ring PET scanners. Images from fPET studies suffer from partial volume errors and residual scatter noise that may cause the cerebral metabolic functional maps to be biased. Gaussian smoothing filters used to denoise the fPET images are suboptimal, as they introduce additional partial volume errors. In this work, a post-processing framework based on a magnetic resonance (MR) Bowsher-like prior was used to improve the spatial and temporal signal to noise characteristics of the fPET images. The performance of the MR guided method was compared with conventional Gaussian filtering using both simulated and *in vivo* task fPET datasets. The results demonstrate that the MR guided fPET framework reduces the partial volume errors, enhances the sensitivity of identifying brain activation, and improves the anatomical accuracy for mapping changes of brain metabolism in response to a visual stimulation task. The framework extends the use of functional PET to investigate the dynamics of brain metabolic responses for faster presentation of brain activation tasks, and for applications in low dose PET imaging.

## 1 Introduction

Brain imaging using positron emission tomography (PET) can provide unique insights into brain function in both healthy individuals and individuals with neuropathological conditions (Nasrallah & Dubroff, 2013). [18F]-fluorodeoxyglucose (FDG)-PET imaging has long been a proxy for regional and global brain metabolism, as glucose uptake is closely correlated with the underlying neuronal activity (Figley & Stroman, 2011; Phelps et al., 1979; Reivich et al., 1985). Conventional static FDG-PET based on a bolus injection of the radiotracer provides a snapshot of glucose metabolism over a long time-window (equal to the scan duration, usually 10-30 minutes). Dynamic PET imaging using a bolus administration of radiotracer provides an opportunity to model tracer kinetics in the brain. However, conventional bolus injection FDG PET scans are not sensitive to cerebral metabolic changes over an extended time duration due to lack of sustained supply of FDG to the brain (Villien et al., 2014). To circumvent this problem, Villien et al. (2014) used a continuous infusion radiotracer infusion approach, together with dynamic PET scanning, to achieve enhanced sensitivity for tracking dynamic radiotracer uptake. This constant infusion approach using FDG was labelled ‘functional’ PET (fPET), to highlight similarities to the functional magnetic resonance imaging (fMRI) technique. Subsequent research using fPET methodology has shown promising results for isolating brain functional areas during external tasks and at rest (Hahn et al., 2016, 2018; Jamadar et al., 2019; Li et al., 2020; Rischka et al., 2018). Despite great improvement in temporal resolution in comparison to traditional approaches, the temporal resolution of fPET remains substantially lower than that of fMRI, which is in the order of seconds or even sub seconds. The current temporal resolution of fPET (around 20-60 seconds) limits the opportunity to use fPET for detailed investigations of brain metabolic responses to rapidly switching tasks and brain stimulation paradigms.

Analysis of fPET data is challenging because of the relatively poor signal to noise ratio (SNR) and partial volume errors in the reconstructed PET images (Z Chen et al., 2018). Recent work has improved the SNR in fPET by applying a combined bolus and continuous infusion of radiotracer during experiments (Jamadar et al., 2019; Rischka et al., 2018). However, the statistical power of these experimental approaches is still relatively low when compared with fMRI. To mitigate this issue, spatial smoothing of the reconstructed PET images is performed prior to functional analysis of the brain using techniques such as independent component analysis (ICA). Gaussian smoothing is widely used as a post-reconstruction spatial and temporal smoothing operation for functional neuroimaging analyses (Zikuan Chen & Calhoun, 2018; Hahn et al., 2018; Jamadar et al., 2019; Pignat et al., 2013; Villien et al., 2014). However, the Gaussian kernel acts as a low-pass filter, and therefore, further worsens the partial volume errors in fPET images; this can cause errors in the localisation and quantification of brain functional activations and at high-temporal resolution fPET imaging. MRI-based PET reconstruction methods have shown substantial improvement in PET image quality compared to conventional methods (Z Chen et al., 2018; V.P. Sudarshan, Chen, & Awate, 2018; Viswanath P. Sudarshan, Egan, Chen, & Awate, 2020). For instance, several studies have explored post-reconstruction PET image enhancement using anatomical information from structural MRI (Bousse et al., 2012; Hutton et al., 2013; Schramm et al., 2018) to perform partial volume correction and image deblurring (Dutta, Leahy, & Li, 2013; Song et al., 2019). The Bayesian formulation of MRI assisted PET denoising can be interpreted as a guided filter to address the PET denoising and partial volume error problems, by modelling the statistical dependencies across the PET and MRI images in order to delineate tissue boundaries.

Loeb et al. (2015) proposed a variant of the well-known Bowsher prior (Bowsher et al., 1996), modelled as prior information in the reconstruction process. The Bowsher prior, in principle, is a weighted Markov random field (MRF) model which promotes delineation of PET image voxels that are dissimilar according to the intensities in the spatially co-registered MRI image. The weights are computed based on a similarity metric (e.g. absolute difference) evaluated on the structural image. Subsequently, Schramm et al. (2018) proposed an asymmetrical variant of the original Bowsher prior and demonstrated that the asymmetrical version yielded PET image reconstruction with improved bias-variance trade-off in comparison to other image gradient-based priors such as parallel level sets (Ehrhardt et al., 2015) and compared to the originally proposed Bowsher prior.

In the current study, we hypothesized that accurate identification of brain metabolic activations could be obtained by filtering the fPET images using knowledge from the anatomical MRI image. The anatomical information was modelled as an MRF prior within a Bayesian framework to restore the fPET signal. The anatomical prior was expected to improve the identification of independent signal components from the fPET data by improving the spatial and temporal SNR and reducing partial volume errors. The formulation of the prior model in this paper differed from the one proposed in (Loeb, Navab, & Ziegler, 2015; Schramm et al., 2018), in that it used a location-dependent smoothly-decaying function incorporating patch-level differences (as opposed to voxel-level differences) to estimate the weights within the neighbourhood of a voxel. The method is henceforth referred to as *MRI-MRF prior* and was validated using both simulated *in vivo* visual task fPET datasets. The accuracy of the method was compared with conventional smoothing methods at both the subject and group level ICA, and the *in vivo* fPET dynamic data were downsampled to verify the robustness of the proposed method in response to reduced task stimulation durations.

## 2 Methods and Experiments

### 2.1 Theory

Let 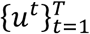 represent the dynamic sequence of *T* fPET images, each containing *N* voxels, reconstructed using model-based iterative methods such as maximum likelihood expectation-maximization (MLEM) (Shepp and Vardi 1982). To perform spatial ICA, we construct a spatiotemporal data matrix, *Y*, using 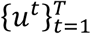, such that the dimension of *Y* is *T* x *N*. ICA models *Y* as a linear combination of the underlying independent components: *Y* = *AS*, where *S* contains the independent components and *A* is the mixing matrix. In the context of PET imaging, the measured PET data is affected by the blurring matrix, *H* (Bousse et al., 2012; Zhu, Gao, & Rahmim, 2019), and the ICA model becomes

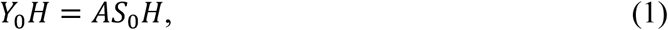

where *Y*_0_ represents the spatiotemporal matrix constructed from the true PET signals, and *S*_0_ models the true underlying independent components of *Y*_0_. The matrix, *H*, acting on the spatial dimension, models the partial volume errors in PET measurements, and hence, the resultant independent components though the mixing operation, *A*.

The goal of fPET data analysis is to identify *S*_0_ from Equation (1). Image denoising in the spatial domain is an important pre-processing step prior to application of the ICA algorithm. The characteristics of an ideal filter for estimation of the source components, *S*_0_, would be to recover the signal without compromising the independence of the underlying true components. Typically, a Gaussian smoothing filter with a suitable width, specified by its full width at half maximum (FWHM) is used to reduce spatial noise, for example, during fMRI data analysis. However, performing a Gaussian smoothing can introduce additional bias in fPET images and the corresponding independent components, due to worsening of the partial volume errors (Zikuan Chen & Calhoun, 2018; Pignat et al., 2013). Hence, this work proposes an MRI guided filtering scheme that can perform (i) denoising, as well as (ii) partial volume correction, to provide an improved estimation of the underlying source components, *S*_0_.

Given the sequence of fPET images, 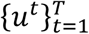, and the fixed MRI image, *v*, of the subject, the post-reconstruction restored fPET image, 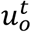, can be obtained by solving the following optimization problem independently for each frame:

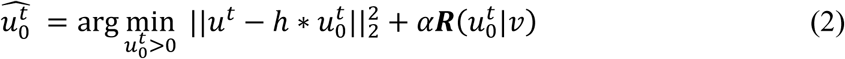

Here ***R***(·) represents the MRI-guided MRF (MRI-MRF) regularization function which incorporates the anatomical information from MRI image, *v*. The kernel function *h* models PSF for current estimate of the image, 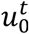. The parameter *α* determines the strength of the regularization, ***R***(·). The formulation in Equation (1) is generic and allows incorporation of arbitrary prior models that enforce certain type of regularity, e.g. piecewise smoothness, on the fPET images. In this work, we model ***R***(·) as a modified version of the asymmetrical Bowsher prior presented by Schramm et al. (2018). Specifically, ***R***(·) is modelled as a weighted quadratic MRF function defined as, ***R***(*u*|*v*) = ∑_*i*∈*I*_ ∑_*j*∈*I*_ *w*_*ij*_(*u*_*i*_ − *u*_*j*_)^2^. Here the weights *w*_*ij*_ are computed based on the intensity values from the co-registered MRI image, *v*, as

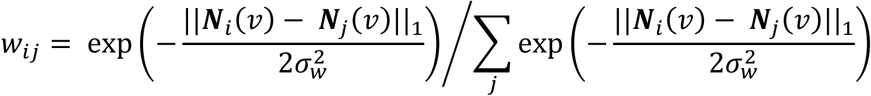

where the operator ***N*_*i*_**(·) extracts a vectorized isotropic 3D patch of volume *L*^3^ mm^3^ centred around voxel *i*, and the parameter σ_*w*_ determines the spatial pattern of weights within the patch in the neighbourhood of voxel *i*. The strategy of determining the weights *w*_*ij*_ in the MRF-based regularization term by relying on patch-difference norms has been used within the literature on patch-based denoising methods, first proposed on natural images in the works of (Awate & Whitaker, 2005a; Buades, Coll, & Morel, 2005) and on MRI images in the works of (Awate & Whitaker, 2005b, 2007; Coupé, Manjón, Robles, & Collins, 2012). While a high value of σ_*w*_ leads to weights that are similar for all the neighbouring voxels, a low value of σ_*w*_ assigns higher weights to a few selected voxels in the neighbourhood. The latter scenario leads to an extension of the strategy in the asymmetric Bowsher prior (Schramm et al., 2018) that (i) enforces neighbourhood weights to be binary (1 or 0) and (ii) design weights only based on voxel-intensity differences (instead of patch differences). Our proposed strategy of using patch-based differences can provide additional robustness to noise and artefacts while leading to better structure preservation, in ways that are similar to those studied in general image denoising (Milanfar, 2012). While iterative denoising algorithms, as in (Awate & Whitaker, 2005a, 2006), offer algorithms for data-driven tuning of the parameter σ_*w*_ to improve performance, in the application in this manuscript, where the weights only need to be precomputed once, we tune the parameter σ_*w*_ based on validation on simulated data.

### 2.2 Data and experiments

Both simulated and *in vivo* fPET and MRI data were used to validate the performance of the MRI-MRF prior. For comparison, the MRI-MRF prior processed fPET images were compared with those obtained using Gaussian smoothing with varying kernel sizes (specified by FWHM).

#### 2.2.1 Simulated experiments and data

Continuous infusion of FDG PET activity was simulated for 60 minutes using a two-tissue compartment model involving the three kinetic parameters *k*_1_, *k*_2_ and *k*_3_ and a fitted arterial input function from intravenous blood samples collected from our previous *in vivo* experimental data (Jamadar et al., 2019). The simulated FDG activity was corrected by the blood partition fraction and haematocrit using the same procedure as in our previous work (Li et al., 2020). Brain regions were segmented into grey matter, white matter and the occipital cortices using the MNI structural atlas (Mazziotta et al., 2001) using FSL (Diedrichsen, Balsters, Flavell, Cussans, & Ramnani, 2009). The MRI and PET images were simulated with an isotropic spatial resolution of 2 mm in the MNI space. The regional specific metabolic kinetic parameters used for the simulated dataset were *k*_1_ = 0.101, *k*_2_ = 0.071, *k*_3_ = 0.042 for grey matter and *k*_1_ = 0.047, *k*_2_ = 0.070, *k*_3_ = 0.035 for white matter, respectively (Lucignani et al., 1993). A visual task stimulus was simulated between 20 to 30 minutes in the visual cortex region similar to the *in vivo* experimental paradigm. During the visual stimulation period, the parameter *k*_3_ in the occipital cortex was simulated to have a 20% increment.

The tomographic iterative GPU-based reconstruction toolbox (TIGRE) was used for PET image reconstruction (Biguri, Dosanjh, Hancock, & Soleimani, 2016). The PET images were forward projected, and Poisson noise was applied in the measurement space, to generate a high-dose dataset. Subsequently, we simulated dynamic low-dose PET data using the Poisson thinning approach (Kim et al., 2018) such that the low-dose data had a dose reduction factor (DRF) of 100 compared to that of the high-dose data. The PET sinogram data were further smoothed in the sinogram space using a Gaussian filter with kernel size 2.35 mm to simulate the partial volume effect. Finally, the MLEM algorithm was used to reconstruct the PET images for the low and high dose datasets.

The reconstructed PET images, 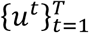, were registered to the corresponding MRI image, *v*. The Bayesian optimization problem with the MRI-MRF prior in Equation (2) was solved using limited memory BFGS method (L-BFGS) (Byrd, Lu, Nocedal, & Zhu, 1995), with positivity constraints. The ICA-specific pre-processing steps including spatial normalization and dimensionality reduction were performed as described in detail by Li et al. (2020) on the post-reconstruction smoothed images. In this work, we performed both subject-level and group-level ICA on fPET data. For group analysis, the spatiotemporal matrix from each subject was concatenated along the temporal dimension before the application of ICA. The pre-processed data, which was an estimate of *Y*_0_, was then decomposed using an ICA unmixing algorithm in the FastICA toolbox (A. Hyvarinen & E. Oja, 2000; Hyvärinen & Oja, 1997).

#### 2.2.2 *In vivo* experiments and data

A cohort of five healthy volunteers were scanned for a visual task stimulus experiment using a 3T Siemens Biograph mMR (Siemens Healthiness, Erlangen, Germany) PET-MRI scanner, approved by the institute human ethics committee. The overall stimulation protocol consisted of three visual stimulation periods consisting of alternating periods of rest and task blocks. A detailed description of the experiment is provided in our earlier work in (Jamadar et al., 2019). The subjects rested for a period of 20 minutes to allow sufficient FDG accumulation in the brain, during which structural MRI scans were acquired. Following this, the subjects viewed a circular flickering checkerboard stimulus for 10 minutes. The checkerboard was retained for a period of 120 seconds and subsequently, an intermittent 32 seconds on and 16 seconds off design was employed. Following the first task stimulation, which involved 3 blocks: rest, task, and rest, two other stimulation experiments, using the full checkerboard visual, were carried out. We used the PET data acquired during the first full checkerboard. Hence, the PET data for each subject was of 30-minute duration, including 10 minutes resting before the stimulation, 10 minutes of a full checkboard stimulation followed by another 10 minutes of rest (Figure 3 (a)). The average dose of FDG given to each subject was 95.9±5.9 MBq which was infused at a constant rate of 36mL/hr over the 90-minute duration.

**Figure 1.**
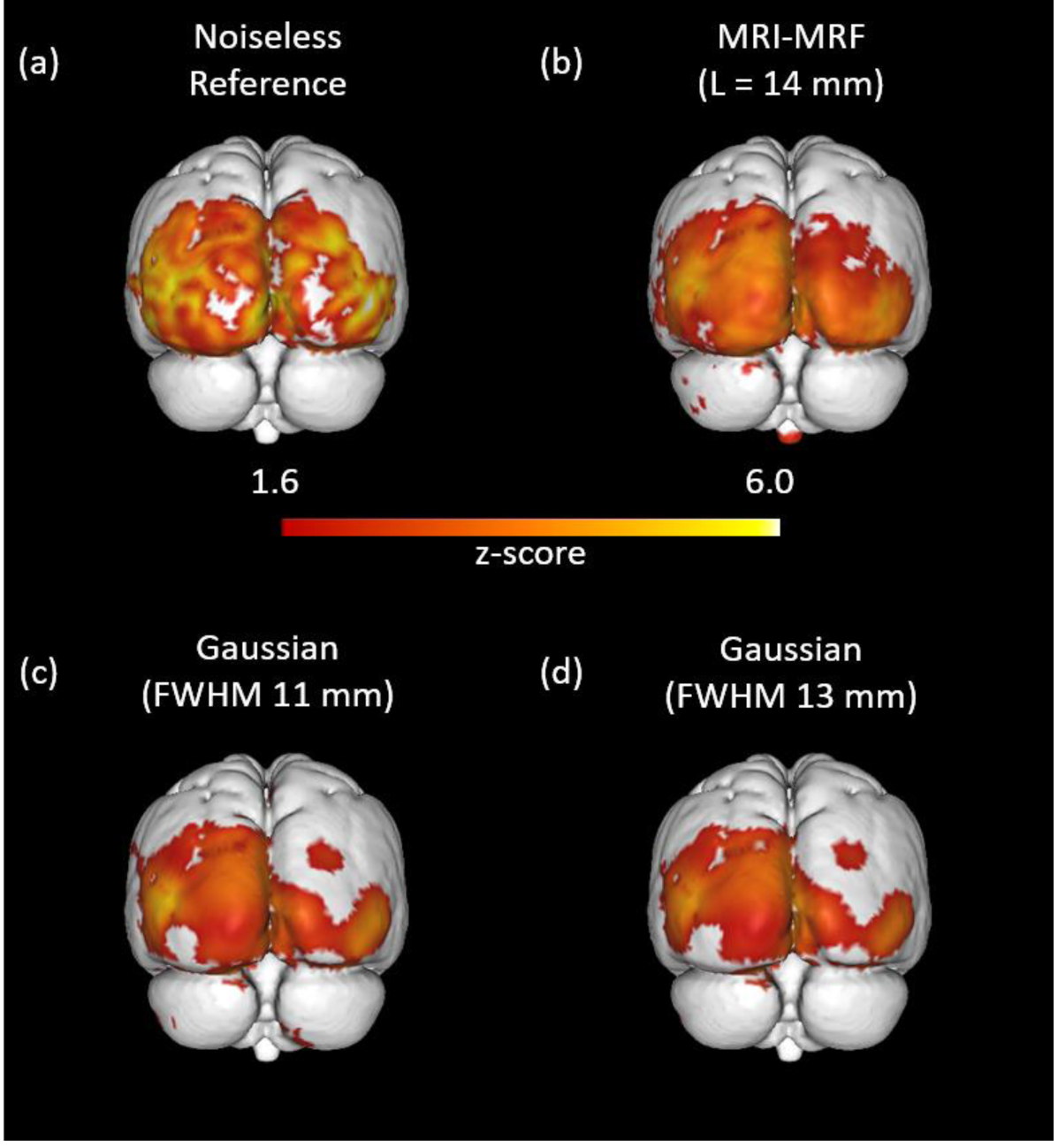
Comparison of brain activation maps using the simulated data. Visualization of activation in the visual cortex using ICA on noiseless fPET images (a), MRI-MRF prior (b), Gaussian smoothing with FWHM 11 mm (c), and FWHM 13 mm (d).

**Figure 2.**
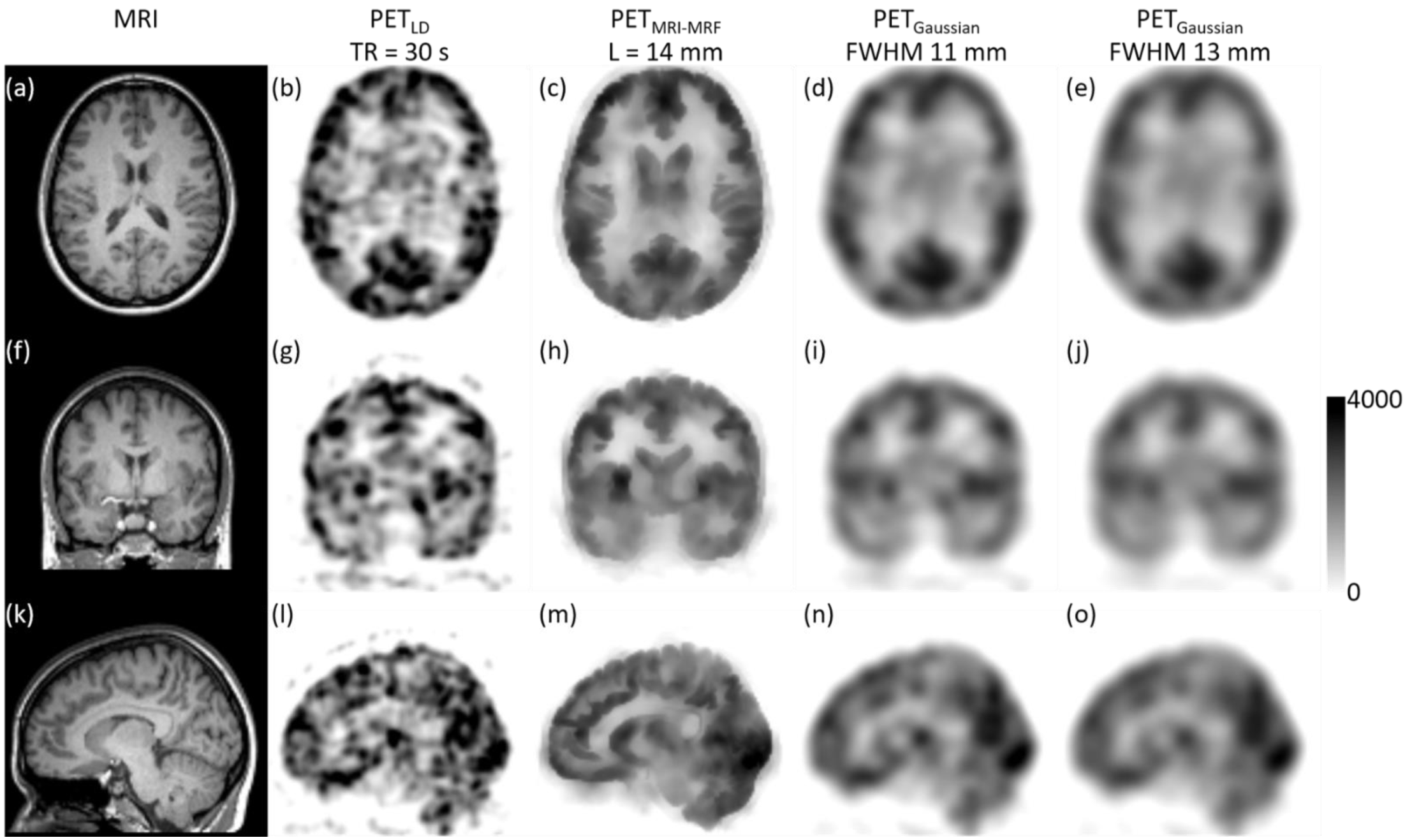
Assessment of the post-reconstruction smoothed fPET images with binning time of Tbin = 30 s. The subject’s MRI image (a); the vendor reconstructed PET image (b); the filtered image using the MRI-MRF prior (L = 14 mm) (c); and using Gaussian kernels with FWHM (d) 11 mm, (e) 13 mm.

**Figure 3.**
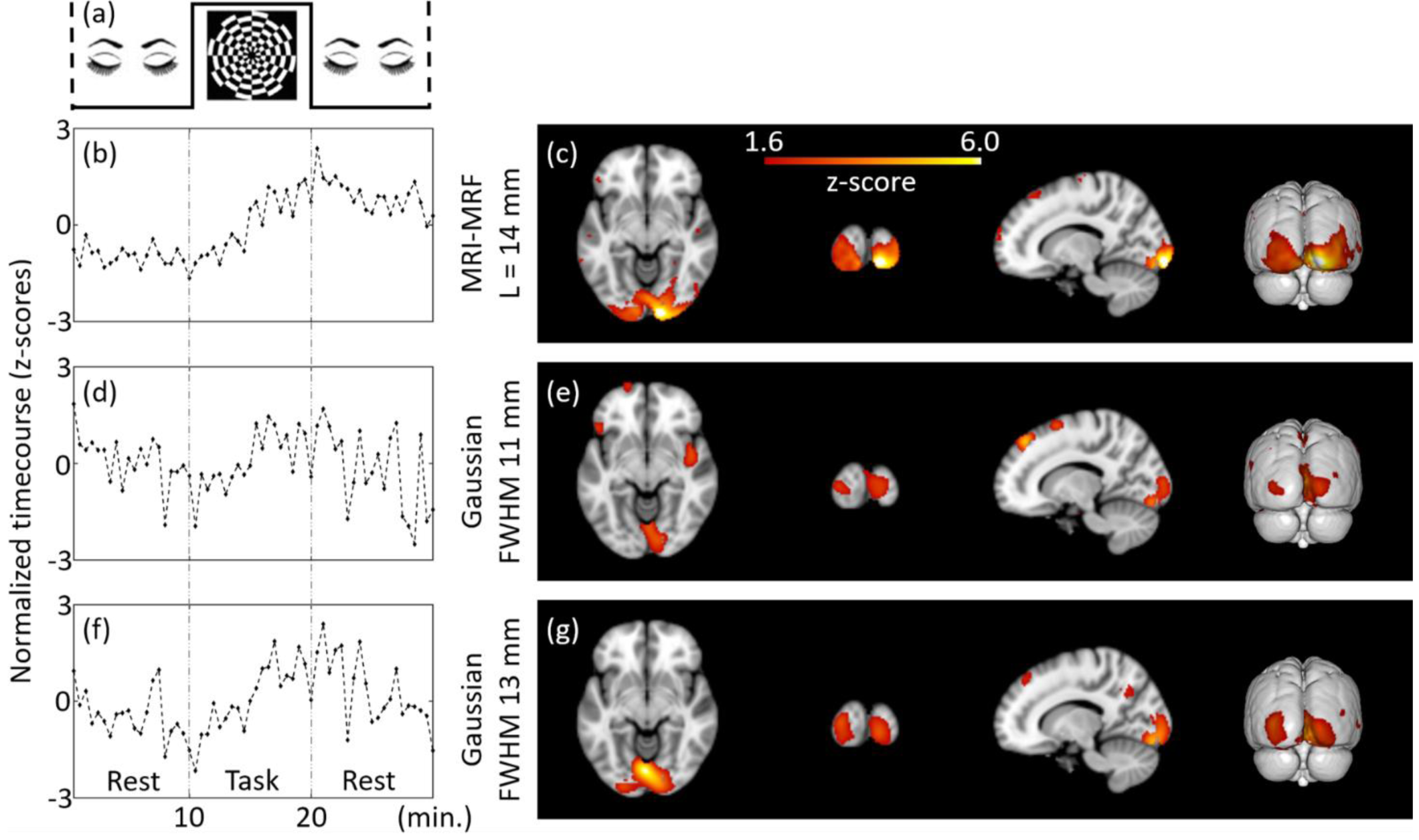
Subject-level (representative) estimation of brain activations using ICA for Tbin = 30 s and DF = 1 at MNI co-ordinate (14, -94, -8). The independent components estimated from the filtered fPET images using different schemes are provided. The task paradigm is shown in (a). ICA maps and timecourses: top to bottom: MRI-MRF prior with L = 14 mm (b) and (c), Gaussian smoothing with FWHM = 11 mm (d) and (e), Gaussian smoothing with FWHM 13 mm (f) and (g).

We reconstructed PET images from the list-mode data using two different values for the temporal bin-width (Tbin) of (i) 30 seconds for the low-dose PET images, and (ii) 3 minutes for high-dose PET images. The average dose for the corresponding low dose fPET images across the group of subjects was calculated to be 7.5 kBq/kg/frame. The PET data was corrected for attenuation using a pseudo-computed tomography (pCT) map (Baran et al., 2018; Burgos et al., 2013). The corrected PET data sinograms were reconstructed using ordered subsets expectation maximization (OSEM) algorithm with 3 iterations and 21 subsets along with point spread function modelling. The PET 3D volumes were reconstructed with voxel sizes of 3 x 3 x 2.03 mm^3^. For standard analysis, all the images were registered to the MNI-152 template. The high-dose PET images from the 3-minute binned data were used to register the low-dose PET images with the T1 weighted MRI (acquired at 1 mm^3^ isotropic resolution) for each subject using ANTS (Avants et al., 2011).

We also undertook a comparison of the performance of the MRI-MRF and Gaussian filtering schemes when the duration of the task and resting blocks was reduced. This analysis was carried out by downsampling the total number of low-dose fPET images reconstructed from the list mode data. Functional PET analyses were computed at both the subject-level and group-level for downsampling factors (DF) of 2 and 3 to simulate fPET images of duration 30 secs but acquired at 1:00 minute and 1:40 minute intervals, respectively. The downsampled PET images correspond to reduced task duration with a lower number of temporal frames.

#### 2.2.3 Optimal kernel width selection

The optimal kernel sizes for the Gaussian low pass filter and the MRI-MRF prior, for processing the *in vivo* data were selected and validated using simulated data. We optimized the parameters to achieve high sensitivity without substantial loss of specificity using ICA computed activation maps. For computing the sensitivity and specificity values, the region of interest (ROI), occipital cortex, was obtained using the segmentation procedure as described in Section 2.2.1. The sensitivity and specificity performance metrics defined as follows:

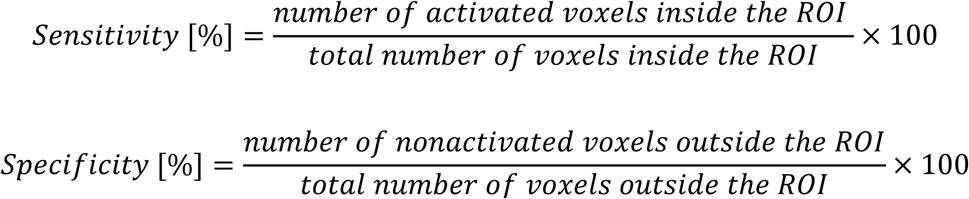

provide a quantitative assessment of the activation maps (z-score map) obtained from the different filtering operations. The metrics were computed by considering a voxel as activated if |*z*| ≥ 1.6 and |*z*| ≥ 2.1, for subject-level and group-level analysis respectively (Li et al., 2020). The parameter search-space for the MRI-MRF prior, includes varying values of the regularization parameter, patch length (*α, L* respectively). On the other hand, for the Gaussian kernel, we varied the FWHM parameter which in turn determines the kernel size.

## 3 Results

### 3.1 Results for simulated data

Table 1 compares the sensitivity and specificity for both denoising schemes at different parameter configurations. For the MRI-MRF prior, the patch-length was varied from 10 mm to 18 mm which represented a varying patch size of 5 to 9 voxels in each direction, respectively. In the case of Gaussian filtering, the kernel size was determined by the full width at half maximum of the Gaussian function. The FWHM for Gaussian varied from 11 mm to 15 mm. It is to be noted that while parameter *L* (for the MRI-MRF prior) represents the width of the entire kernel, FWHM (for the Gaussian filter) represents approximately half of the kernel-width. The parameter range chosen for the Gaussian smoothing is consistent with the Gaussian kernel widths used in the prior work (Li et al. 2020). The sensitivity values for the MRI-MRF processed image are dramatically higher than that of the Gaussian smoothed images, whereas the specificity values are comparable between the two methods. For the fPET data analysis, a patch-length of 14 mm was chosen for the MRI-MRF prior. However, both the Gaussian kernels with FWHM 11 mm and 13 mm show similar sensitivity and specificity values. Therefore, the analysis using the Gaussian-filtered *in vivo* fPET data was undertaken using both the 11 mm and 13 mm FWHM filters.

**Table 1.**
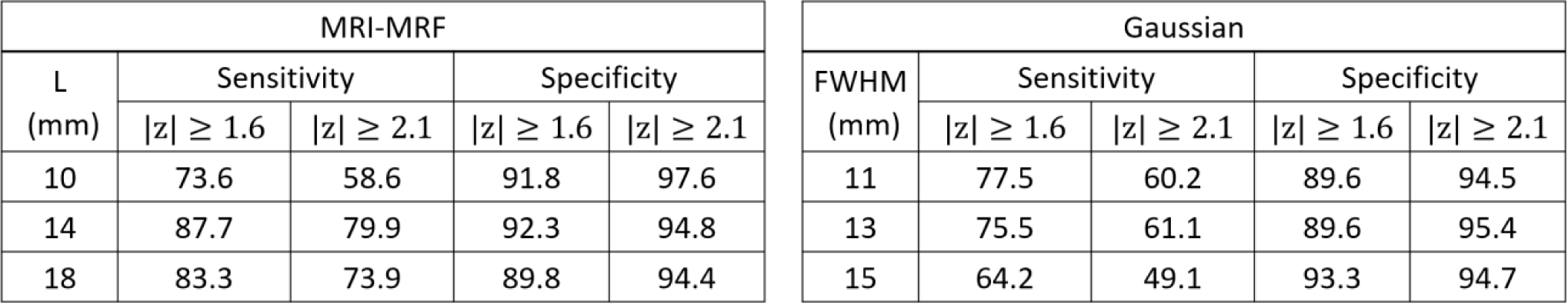
Comparison of sensitivity and specificity of MRI-MRF and Gaussian smoothing filters. The sensitivity and specificity values at two different z-score threshold values are provided.

Figure 1 shows the visual task specific activation for the reference noiseless fPET images, and for the three denoising schemes using the optimal parameters chosen from Table 1. The ICA activation map obtained using the noiseless images serves as the reference map (Figure 1 (a)). The activation map obtained from post-reconstruction filtered fPET images using the MRI-MRF prior (Figure 1 (b)) was closest to the reference activation map in the visual cortex. On the other hand, the activation maps obtained using Gaussian smoothing with both FWHMs yield suboptimal activation maps in the visual cortex with asymmetrical patterns.

### 3.2. Results for *in vivo* data

The results reported in this section are for the fPET images reconstructed using the list-mode data binned at Tbin = 30 s, and for DF = 1, 2 and 3.

Figure 2 shows the post-reconstruction filtered fPET images along with the subject’s MRI image (Figure 2, left column) and the corresponding vendor provided low dose fPET image (Figure 2, second column). The denoised image using the MRI-MRF prior (Figure 2, third column) shows superior recovery of PET signals in different regions of the brain while removing substantial amount of noise. Specifically, the white and grey matter boundaries are well delineated, the shape of the ventricles has been recovered (which is not evident in the low dose PET image), and anatomical features in the gyri, sulci and details of the cortical folding (refer Figure 2m) have been restored. On the other hand, the denoised images using both Gaussian kernels (FWHM 11 mm and 13 mm) are heavily blurred and show substantial loss of anatomical details due to the partial volume errors (Figure 2, fourth and fifth columns).

Figure 3 shows the results of an individual subject-level fPET analysis obtained using different filtering techniques for a downsampling factor of one (i.e. DF=1, includes all list-mode data). The ICA activation maps corresponding to the visual task component along with the normalized timecourses (representing the z-scores) are calculated for each filtering method. The component maps for all sections of the brain are provided in the Supplementary material. The ICA timecourse for both Gaussian kernels (Figures 3d & 3f) are noisy and do not closely follow the experimental task paradigm. Moreover, the shape of the region of brain activation does not follow the known anatomical structure of the primary visual cortex but extends into adjacent neuroanatomical structures including the white matter, likely due to large partial volume errors. Conversely, the activation map obtained using the MRI-MRF prior (Figure 3c) shows localized activity near the visual cortex with a significantly higher z-score within the visual cortex compared to both Gaussian kernels. The ICA timecourse for the MRI-MRF prior (Figure 3b) accords more closely with the experimental design with increased uptake during the visual task block. The comparison of the visual task components for the three methods for all brain sections is consistent with these observations (see Supplementary material).

The results for the group-level fPET analyses for the three filtering techniques using the complete list-mode dataset (DF = 1) are shown in Figure 4. A higher z-score range was observed for the group-level analyses compared to the single subject-level analysis. However, in contrast to the subject-level analysis, the timecourses estimated from all methods (Figures 4b, 4d & 4f) at the group level recapitulated the experimental design paradigm. The activation map corresponding to the MRI-MRF prior followed the known neuroanatomical representation of the primary visual cortex and was consistent with the subject-level result. On the other hand, the activation maps using the two Gaussian kernels did not represent activation in the primary visual cortex and demonstrated diffuse cerebral metabolic activity into large adjacent anatomical regions including the white matter.

**Figure 4.**
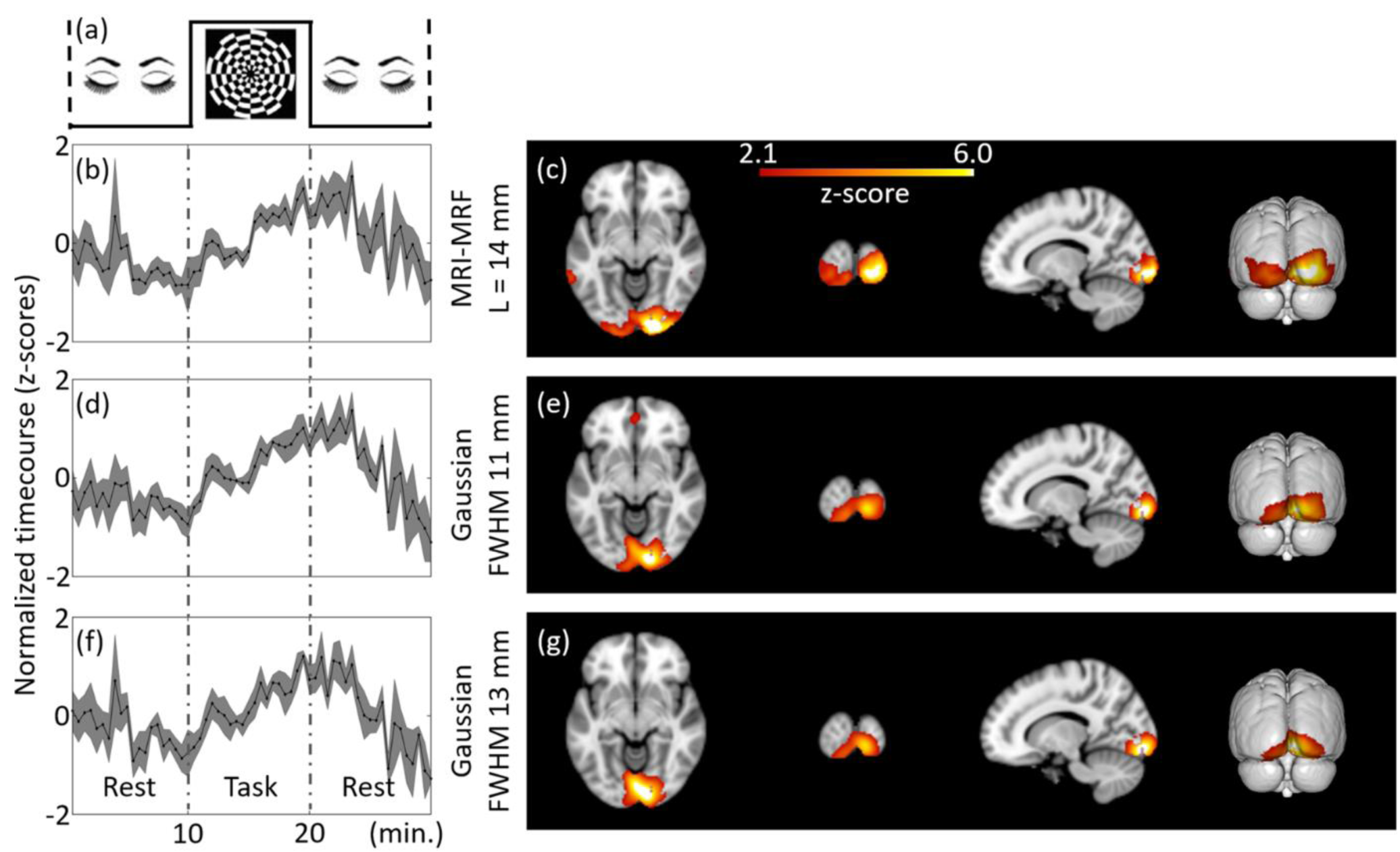
Group-level estimation of brain activations using ICA for Tbin = 30 s and DF = 1 at MNI co-ordinate (14, -94, -8). The independent components estimated from the filtered fPET images using different schemes are provided. The task paradigm followed for the group study is shown in (a). ICA maps and timecourses: top to bottom: MRI-MRF prior with L = 14 mm (b) and (c), Gaussian smoothing with FWHM = 11 mm (d) and (e), Gaussian smoothing with FWHM 13 mm (f and (g)).

Figure 5 shows the subject-level fPET analyses for downsampling factors of two and three (DF = 2 & 3). Timepoints T1 and T2 represent the onset of the task and second resting block in the downsampled task paradigm. Plausible ICA activation maps were not generated using an 11mm FWHM Gaussian kernel for both DFs and therefore no results are included. The ICA timecourses during the task-block for the MRI-MRF filter demonstrated a steadier gradual increase, in agreement with the task paradigm, in comparison to the 13mm FWHM Gaussian kernel for DFs of 2 and 3 respectively (Figures 5a & 5e compared to Figures 5c & 5g respectively). The activation map axial view for DF = 3 did not reveal activation in the left hemisphere as was expected for the visual task (Figure 5h). However, for DF = 2 there was some activation in the left hemisphere visual cortex (Figure 5d) although it was not as widespread as for the fully sampled dataset. On the other hand, the activation maps for the MRI-MRF prior (Figures 5b & 5f) showed spatial congruency across the three DFs, whilst the discrepancy between the z-scores for the MRI-MRF prior and the 13 mm FWHM Gaussian filter was largest for DF =3 compared to DF = 2 and 1.

**Figure 5.**
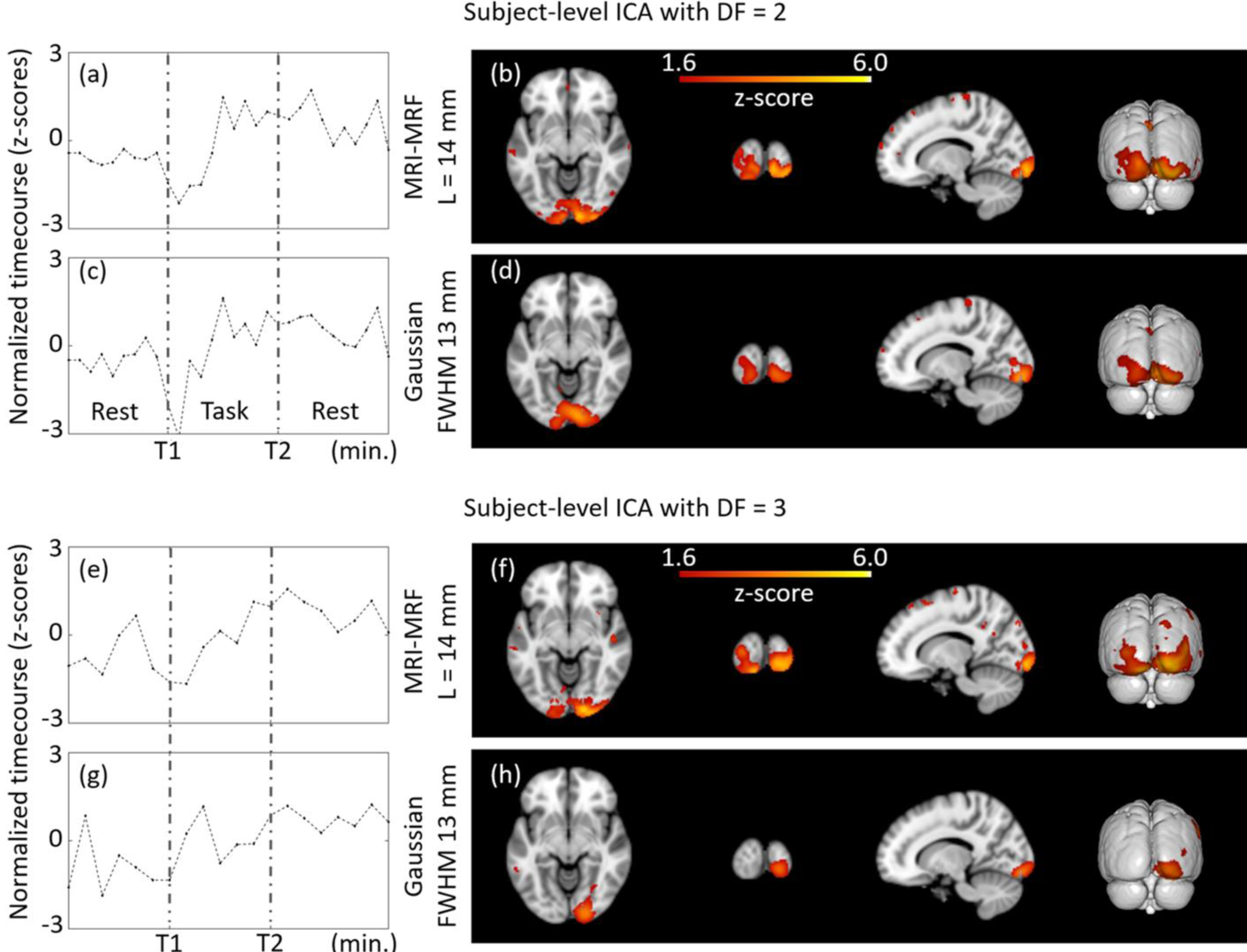
Subject-level (representative) estimation of brain activations using the reduced task and resting blocks with DF = 2 and 3 at MNI co-ordinate (14, -94, -8). The independent components estimated from the filtered fPET images using different schemes are provided. DF = 2: MRI-MRF prior (b) and Gaussian kernel with FWHM 13 mm (d). DF = 3: MRI-MRF prior (f), and Gaussian smoothing with FWHM = 13 mm (h). The T1 and T2 represent the onsets of the task and second resting block, respectively.

Figure 6 shows the group-level fPET analyses at DF = 2 and DF = 3. In contrast to the group-level analysis for the fully sampled dataset where there was little difference between the activation maps estimated by the MRI-MRF method and the Gaussian kernel with FWHM 13 mm (Figure 4), the activation maps estimated for the group-level analyses for DF= 2 and DF = 3 showed marked differences. For both DF = 2 and 3, the ICA timecourses for the MRI-MRF prior (Figures 6a & 6e) showed agreement with the task experimental design with higher z-scores than for the 13mm FWHM Gaussian filter timecourses (Figures 6c & 6g). The activation maps show that while the MRI-MRF prior was able to resolve brain activation that was consistent with activation of the visual cortex (Figures 6b & 6f), at both DF = 2 and 3 the 13mm FWHM Gaussian filter was unable to resolve extended activation throughout the primary visual cortex (Figures 6d & 6h) with no activation in the left hemisphere for DF = 3 (Figure 6h). Conversely, the activation maps for the MRI-MRF prior were congruent across the subject-level and group-level analyses although greater consistency in the right hemisphere.

**Figure 6.**
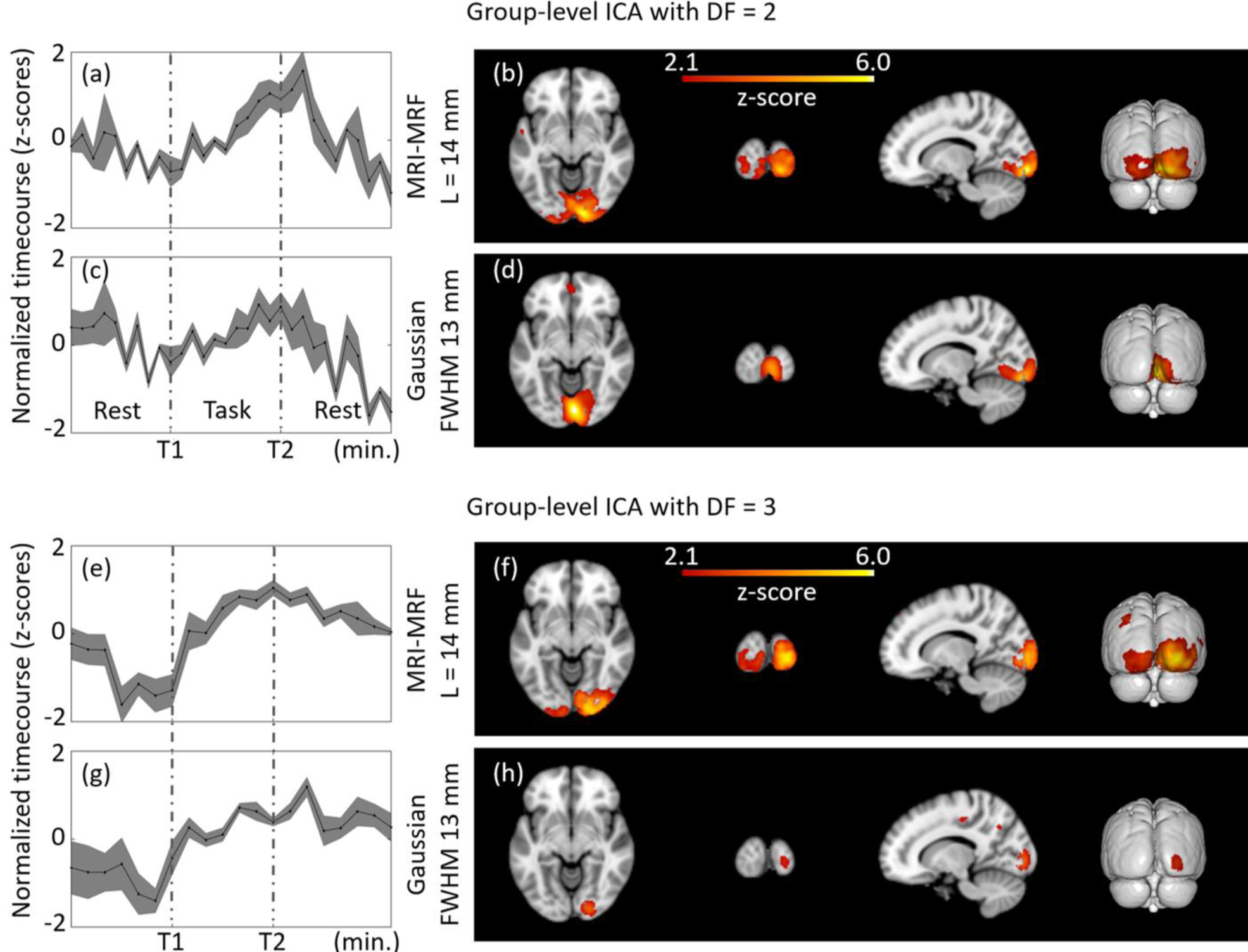
Group-level estimation of brain activations using the reduced task and resting blocks with DF = 2 and 3 at MNI co-ordinate (14, -94, -8). The ICA components estimated from the filtered fPET images using different schemes are provided. DF = 2: MRI-MRF prior (b) and Gaussian kernel with FWHM 13 mm (d). DF = 3: MRI-MRF prior (f), and Gaussian smoothing with FWHM = 13 mm (h).

## 4 Discussion

We have proposed an MRI-assisted fPET processing framework for the analysis of task-related metabolic changes in the brain using high temporal resolution fPET data and for low-dose fPET brain mapping applications. We investigated the effect of using the anatomical information from a subject’s MRI to denoise the fPET dataset to reduce the partial volume error in the PET images in order to increase the sensitivity of the ICA analysis. The PET image restoration problem was posed as a solution to a Bayesian optimization problem which was solved using L-BFGS due to its greater computational efficiency compared to gradient-descent based optimization techniques.

This study compared the post-reconstruction filtered images from the MRI-MRF method and Gaussian smoothing with varying kernel sizes as well as the ICA activation maps from the fPET dataset using a visual stimulation task. Visual assessment of the post-reconstruction smoothed images showed that the MRI-MRF processed PET images recovered many features which were not readily observed in the conventional low dose PET images. The MRI-MRF filtered PET images revealed localized tracer uptake in the sub-cortical nuclei adjacent to the lateral ventricles (e.g. Figure 2c) whereas little or no uptake was apparent in the comparable low-dose and Gaussian denoised PET images. Furthermore, the level of Gaussian smoothing required to obtain plausible activations in the visual cortex rendered the fPET image hard to interpret visually as there was a substantial loss of features. The MRI-MRF method provides a balance between visual interpretability of the fPET images together with improved resolution and sensitivity for functional analysis using ICA.

The task-based experimental design paradigm enabled meaningful comparison of the ICA timecourses obtained using the two filtering techniques, by inspection of the FDG uptake in the visual cortex during the visual stimulation task. The proposed methodology was able to achieve consistent activation maps at both the subject-level and group-level for DF = 1, 2 and 3. However, this was not true for the Gaussian smoothing kernels. Moreover, since the fPET data was acquired for an exogenous task-based stimulus, good correlation between the subject-level and group-level activation maps was expected. In particular, the improved results for the individual subject-level analysis demonstrates the benefit of the MRI-MRF method to enhance single subject-level analysis using low dose high temporal resolution fPET data with reduced task durations.

The study involving downsampling factors that demonstrated that the proposed processing pipeline could detect dynamic brain metabolic increases for visual task stimulation for as short as approximately three minutes. However, this interpretation does assume that the FDG dosage per frame in the fPET images is consistent for the different downsampling factors. In practice this would be achievable experimentally by altering the infusion protocol or slightly increasing the dosage of the radiotracer (Verger & Guedj, 2018). The Gaussian smoothing technique failed to identify task related ICA components for the shorter task durations (i.e. at higher DFs) due to reduced sensitivity.

Unlike fMRI, fPET images suffer from very low SNR and hence the spatial denoising scheme must be carefully chosen to provide an optimal bias-variance trade-off. MRI-guided PET image denoising and deblurring has been extensively reported in the literature (Hutton et al., 2013; Song et al., 2019) with many solutions for post-reconstruction PET image enhancement. However, this paper is the first to demonstrate the effectiveness of the MRI-based spatial denoising technique for dynamic fPET imaging, such that fPET images are both visually interpretable and produce accurate functional maps with improved temporal resolution. The high specificity and sensitivity of the algorithm also enabled single subject-level analyses along with reasonable visualization of the fPET images without loss of anatomical details. Traditional methods such as Gaussian smoothing perform averaging without consideration of the anatomical boundaries and hence the quantitative accuracy of FDG signals is degraded by partial volume errors. This was reflected in the diffuse visual activation maps obtained with the Gaussian filtering. Using edge-preserving denoising techniques such as anisotropic filtering would also yield suboptimal performance because of the poor SNR of the fPET images and the difficulty to distinguish between tissue boundaries and noise.

The formulation of the MRI-MRF prior in this work is generic and allows for modelling of higher-level image features such as dictionary atoms. Nevertheless, the proposed filtering framework is efficient and computationally less expensive in comparison to other patch-based techniques and hence, the framework is easier to adapt for other research and clinical applications.

Although the MRI-MRF prior in this work was applied in the image domain on post-reconstructed fPET images, the same could be applied within the image reconstruction process provided the PET list-mode data was accessible. Research using image restoration techniques in a reconstruction framework generally employ a Poisson noise model for the sinogram data and a system matrix composed of several matrix operations representing the point spread function, forward projection, attenuation correction, scatter correction, and back projection. Our work solved the image restoration problem in the image domain and employed a least-squares-type data term rather than a fixed noise-model in the image space. This was because the noise characteristics of the reconstructed PET images inherently depend on the reconstruction algorithm. For example, noise characteristics during filtered back projection reconstruction depend upon the filter employed, whilst in maximum likelihood expectation maximisation reconstruction and its variants, the noise characteristics depend on the number of iterations as well as the strength of the prior function.

The current work has a number of limitations. One of the limitations is the small sample size. In this work, we show proof of the principle for utilizing anatomical information for fPET data processing. Advanced statistical image restoration models such as joint patch-based techniques and neural networks may further improve the image quality for shorter image acquisition durations and potentially in future approach the temporal resolution offered by fMRI. However, the proposed framework is readily adaptable to use these techniques in the research context although modelling higher statistical dependencies would increase the number of hyperparameters that were required to be tuned. A further limitation is that the MRI-MRF modelled as a Bowsher-like prior may be perceived as a technique that relies excessively on the anatomical modality. Although this may be relatively unimportant or in fact beneficial in the case of tracers like FDG that are widely distributed throughout the brain, this may not be the case for other heterogeneously distributed tracers such as for amyloid PET imaging. More sophisticated image restoration models which maintain a balance between the PET and MRI features for each tracer may need to be incorporated at the cost of more computational time. The use of spatial regularization could be carefully extended to include a temporal smoothing constraint governed by studies in tracer kinetics. A comprehensive study of several MRI-PET joint priors in the context of dynamic functional PET denoising and analytical techniques is an important direction for future studies.

## 5 Conclusion

We have presented a novel MRI-assisted fPET processing framework for functional analysis of fPET data at high temporal resolution and for low doses of radiotracer. Compared to traditional Gaussian smoothing, our approach yields visually interpretable PET images while increasing the sensitivity and anatomical accuracy of activation maps estimated using ICA. Through validation using simulated data, we have demonstrated that the MRI-MRF method is able to accurately estimate visual task related brain activation maps even under poor SNR conditions. The application to *in vivo* fPET data demonstrated that the MRI-MRF prior achieves detection of reduced task durations of approximately three minutes and provides an avenue for further increases in the temporal resolution and sensitivity of both single subject and group-level brain metabolic mapping studies.

## Acknowledgements

The authors are grateful to Richard McIntyre and Alexandra Carey for organizing the scanning protocol. We thank Edwina Orchard, Irene Graafsma and Disha Sasan in the assistance of data collection, and Phillip Ward for useful discussions. We acknowledge the provision of Siemens e7tools used for reconstruction of the PET images.

This work was supported by an Australian Research Council (ARC) Linkage Project (LP170100494) to G.F. Egan, S.D. Jamadar & Z. Chen which includes financial support from Siemens Healthineers. S. Li and Z. Chen are supported by funding from Reignwood Cultural Foundation. G.F. Egan & S.D. Jamadar are supported by the ARC Centre of Excellence for Integrative Brain Function (CE140100007). S.D. Jamadar is supported by a National Health and Medical Research Council of Australia Fellowship (APP1174164). S.P. Awate is supported by the Infrastructure Facility for Advanced Research and Education in Diagnostics grant funded by Department of Biotechnology (DBT), Government of India (BT/INF/22/SP23026/2017).

