## Supplementary Material for "Improved temporal resolution for mapping brain metabolism using functional PET and anatomical MRI knowledge"

### Supplementary Results

#### ICA Results

In the main text, we reported the results of the visual task components obtained both at subject-level and group-level from the proposed MRI-MRF prior and Gaussian smoothing. Here we provide all the comparisons across different slices for both the methods.

Supplementary Figures 1 – 4 compares the visual task components for MRI-MRF prior and the Gaussian smoothing with FWHM 13 mm. At all the DFs, for most of the axial slices, the MRI-MRF prior shows activation which closely follows the anatomy of the visual cortex. In slices where there is increased activation shown by the MRI-MRF prior, results from Gaussian smoothing shows a large blob of activation around the visual cortex but extends to other anatomical regions. At DF = 1 and 2, the Gaussian filter shows activations which do not appear localized in the visual cortex region. At DF = 3, Gaussian smoothing does not provide activation which can be concluded as task related activations in the visual cortex.

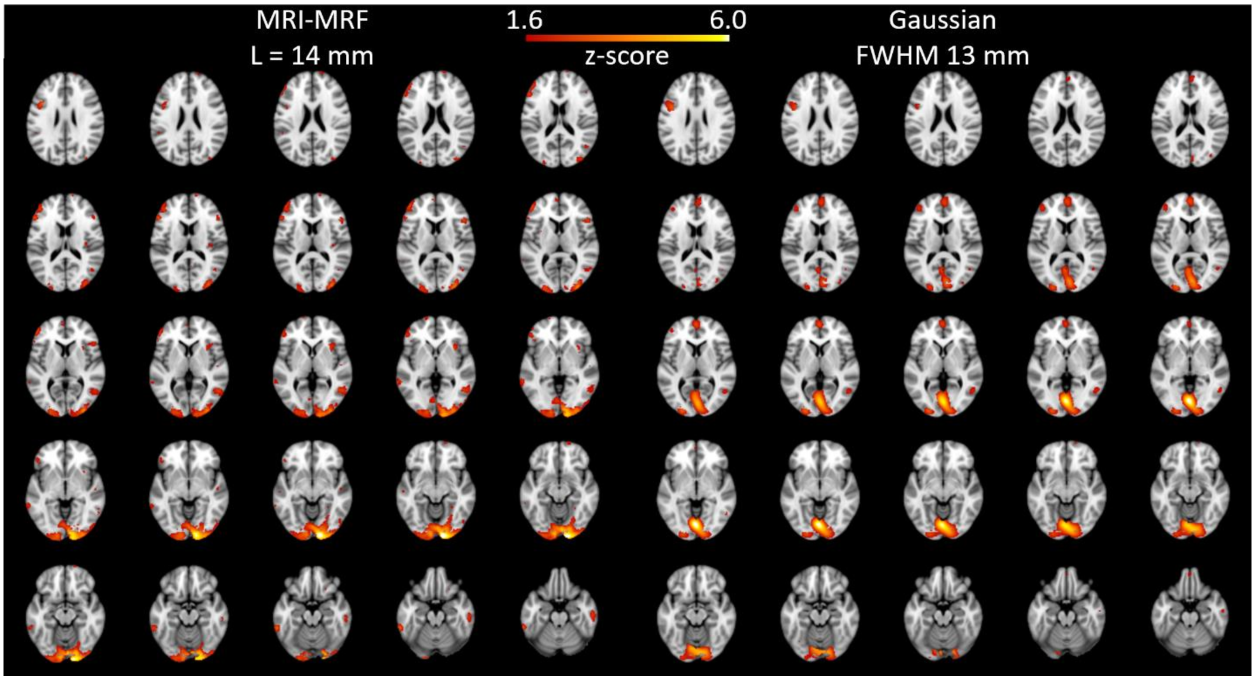


Supplementary Figure 1 Subject-level ICA at DF = 1. Comparison of visual task component across slices for MRI-MRF (left) and Gaussian filter (right) with FWHM 13 mm.

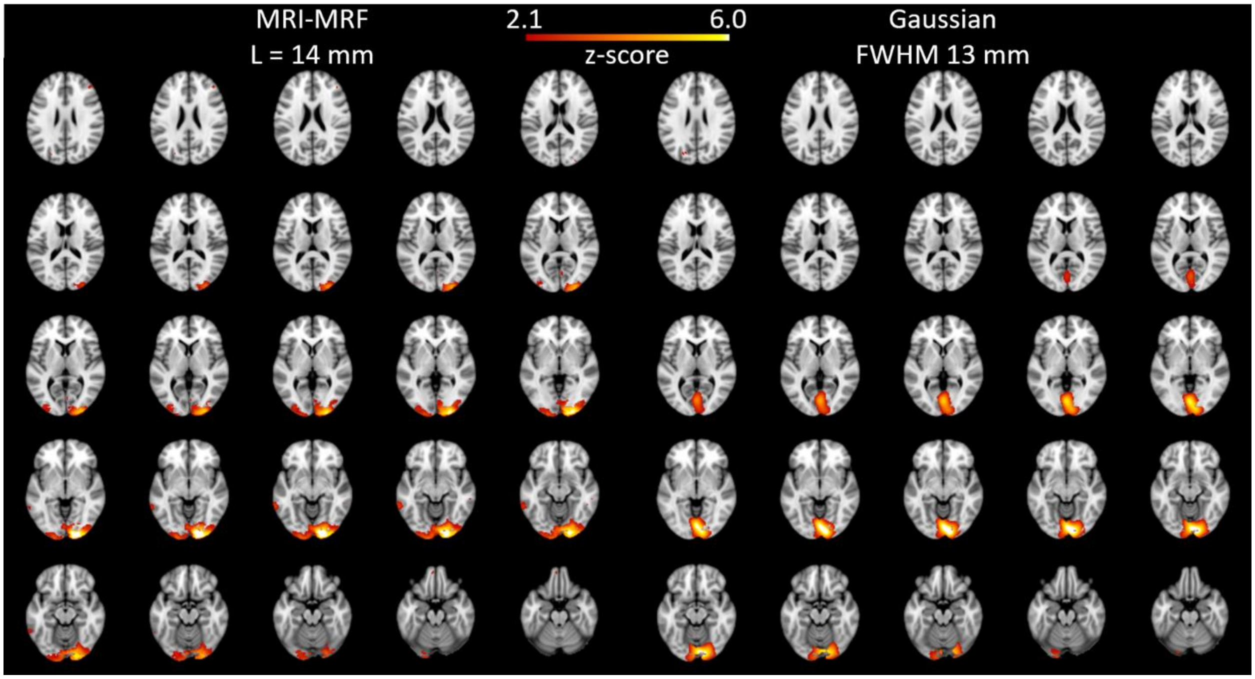


Supplementary Figure 2 Group-level ICA at DF = 1. Comparison of visual task component across slices for MRI-MRF (left) and Gaussian filter (right) with FWHM 13 mm.

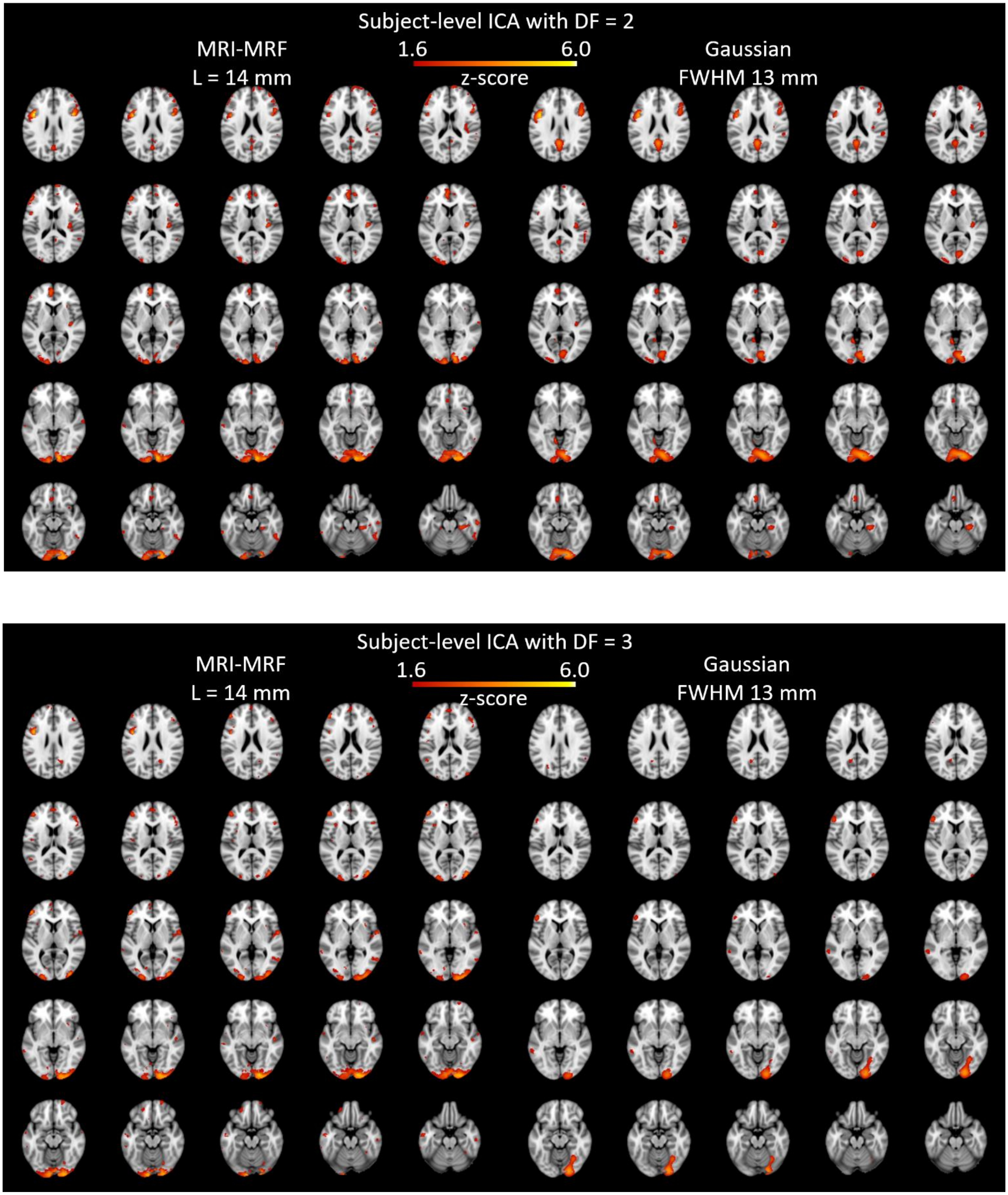


Supplementary Figure 3 Comparison of subject-level ICA between MRI-MRF prior and Gaussian smoothing at DF = 2 (top) and 3 (bottom).

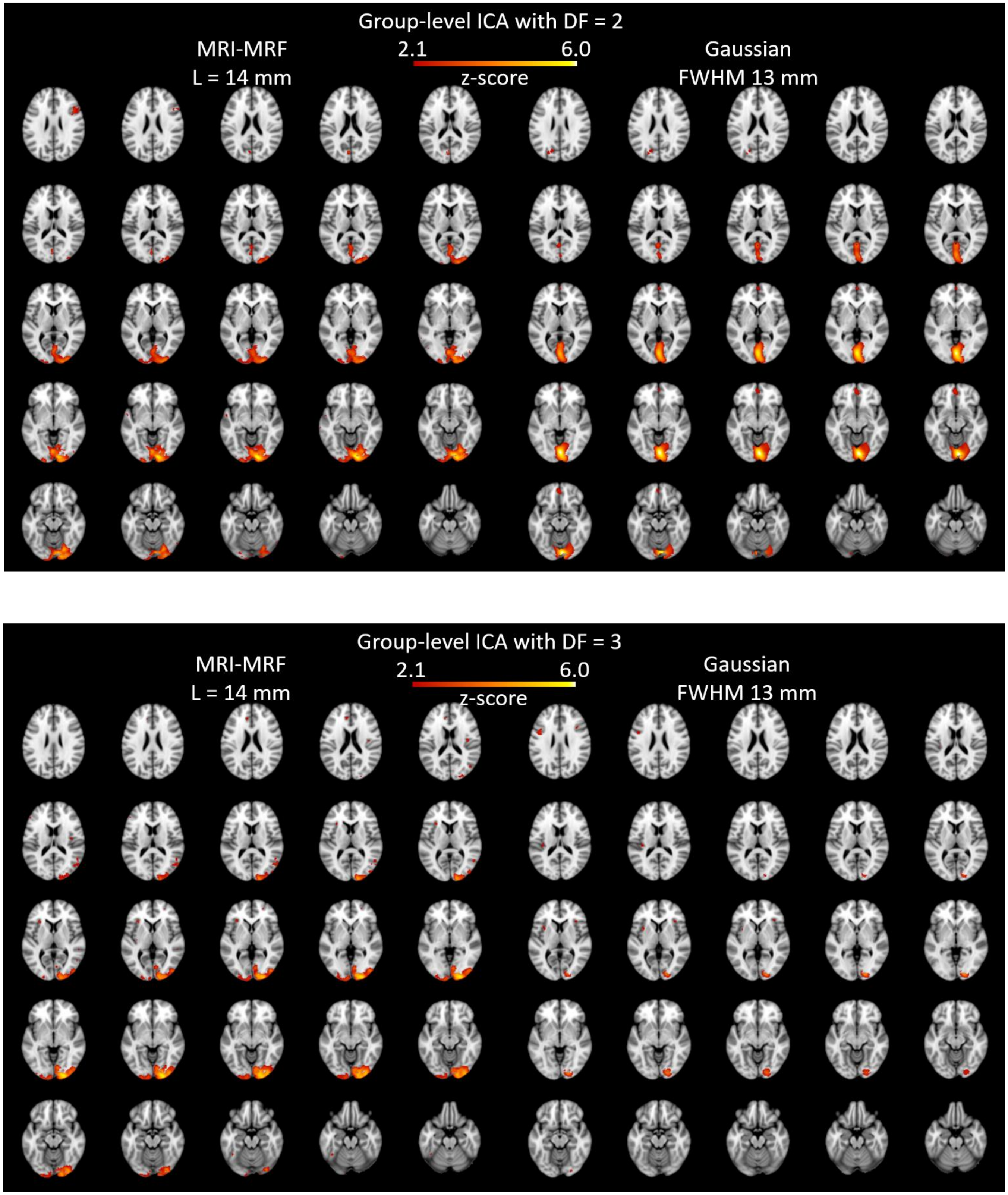


Supplementary Figure 4 Comparison of group-level ICA between MRI-MRF prior and Gaussian smoothing at DF = 2 (top) and 3 (bottom).
